# Chronic adolescent exposure to cannabis in mice leads to sex-biased changes in gene expression networks across brain regions

**DOI:** 10.1101/2021.11.30.470393

**Authors:** Yanning Zuo, Attilio Iemolo, Patricia Montilla-Perez, Hai-Ri Li, Xia Yang, Francesca Telese

## Abstract

During adolescence, frequent and heavy cannabis use can lead to serious adverse health effects and cannabis use disorders (CUD). Rodent models of adolescent exposure to the main psychoactive component of cannabis, delta-9-tetrahydrocannabinol (THC), mimic the behavioral alterations observed in adolescent users. However, the underlying molecular mechanisms remain largely unknown. Here, we treated female and male mice with high doses of THC during early adolescence and assessed their memory and social behaviors in late adolescence. We then profiled the transcriptome of five brain regions involved in cognitive and addiction-related processes. We applied gene coexpression network analysis and identified gene coexpression modules, termed cognitive modules, that simultaneously correlated with THC treatment and memory traits reduced by THC. The cognitive modules were related to endocannabinoid signaling in the female dorsal medial striatum, inflammation in the female ventral tegmental area, and synaptic transmission in the male nucleus accumbens. Moreover, cross-brain region module-module interaction networks uncovered intra- and inter-region molecular circuitries influenced by THC. Lastly, we identified key driver genes of gene networks associated with THC in mice and genetic susceptibility to CUD in humans. This analysis revealed a common regulatory mechanism linked to CUD vulnerability in the nucleus accumbens of females and males, which shared four key drivers (*Hapln4, Kcnc1, Elav12, Zcchc12*). These genes regulate transcriptional subnetworks implicated in addiction processes, synaptic transmission, brain development, and lipid metabolism. Our study provides novel insights into disease mechanisms regulated by adolescent exposure to THC in a sex- and brain region-specific manner.

## Introduction

Cannabis remains the most widely used psychoactive drug worldwide, particularly among adolescents and young adults^1^. Recent data showed that more than one-third of 12th graders in the US used cannabis in the past year, reflecting an overall decline in the perceived risk of regular cannabis use among adolescents^2^. The high prevalence rates of cannabis use among adolescents pose a significant concern as cannabis misuse by adolescents can lead to persistent cognitive impairments in learning, attention, and memory^3–8^. Moreover, the early onset of cannabis use before 16 years of age increases the risk of developing psychiatric disorders, including cannabis use disorders (CUD) ^9,10^. CUD has a strong genetic component (50-70% heritability). Recent large-scale genome-wide association studies (GWAS) began to identify genetic variants associated with CUD^11–15^. They also revealed a genetic correlation of CUD with other mental health traits, including substance abuse, schizophrenia, and risk-taking^11–15^. However, in line with the nature of complex disease traits, common genetic variants associated with lifetime cannabis use can explain only 11% of the phenotypic variance, as revealed by one meta-analysis of eight GWAS^15^. It is possible that other environmental factors, including cannabis exposure during critical developmental periods, might affect molecular networks in critical brain regions and, in turn, influence the development and severity of CUD.

The primary psychoactive component of cannabis is delta-9-tetrahydrocannabinol (THC). The biological effects of THC are mediated mainly by members of the G protein-coupled receptors (GPCR) family, such as cannabinoid receptors 1 (CB1R) and 2 (CB2R)^16^. The cannabinoid receptors, together with endogenous cannabinoids and the enzymes responsible for their biosynthesis and metabolism, constitute the endocannabinoid (eCB) system^17–19^. The eCB system plays a critical role in the maturation of brain circuits during adolescence by regulating excitatory and inhibitory neurotransmission^20^. Moreover, the fluctuations of eCB signaling during adolescence influence the pubertal changes in gonadal hormone secretion^21^. This interaction between eCB signaling and gonadal functions contributes to the emergence of sex-biased behaviors during adolescence, including social, cognitive, and emotional behaviors^22^. Substantial evidence from human or animal model studies has led to the hypothesis that excessive exposure to THC during adolescence may disrupt the physiological function of the eCB system, ultimately leading to sex-specific behavioral abnormalities and increased risk for psychopathology later in life ^5,22–25^.

Despite this knowledge, there is limited data on genes and pathways affected by adolescent exposure to THC in different brain regions of the female and male brains. Two recent studies analyzed gene expression changes following chronic adolescent exposure to THC in rats^26,27^. These studies demonstrated that chronic adolescent exposure to THC alters gene expression networks that are associated with the structural maturation of cortical cells in the prefrontal cortex (PFC) and with reward and stress reactivity in the basolateral amygdala (Amy). However, these analyses were limited to one brain region in male rats ^26,27^. Therefore, more research is needed to dissect brain region-specificity and cross-brain networks perturbed by chronic adolescent exposure to THC in the female and male brains.

In this study, we treated female and male mice with THC during early adolescence and assessed their behavior in late adolescence. We performed RNA sequencing (RNA-seq) on five brain regions involved in cognitive and addiction-related processes, including the prefrontal cortex (PFC), nucleus accumbens (NAc), dorsal medial striatum (DMS), amygdala (Amy), and ventral tegmental area (VTA). We conducted gene coexpression network analysis for each sex, within and between brain regions. Lastly, we performed integrative genomic analyses of coexpression networks altered by THC in mice and human genetic data from GWAS of CUD. This analysis identified genes and pathways involved in THC-mediated behavioral aberrations in mice and linked them to CUD in humans.

## Methods and Materials

Detailed descriptions of experimental design and methods are included in the Supplemental Methods. All methods used in this study have been published recently^28^–^31^. RNA-seq datasets were deposited on the GEO database (accession GSE189821).

## Results

### THC exposure during adolescence impairs cognitive behaviors in a sex-specific manner

To assess the behavioral effects of adolescent exposure to THC, we chronically treated female and male mice with 10mg/kg of THC in early adolescence (5-7 weeks). We assessed object recognition memory, social interaction, and anxiety-like behaviors in late adolescence, two weeks after the last THC administration (Fig 1A).

**Figure 1:**
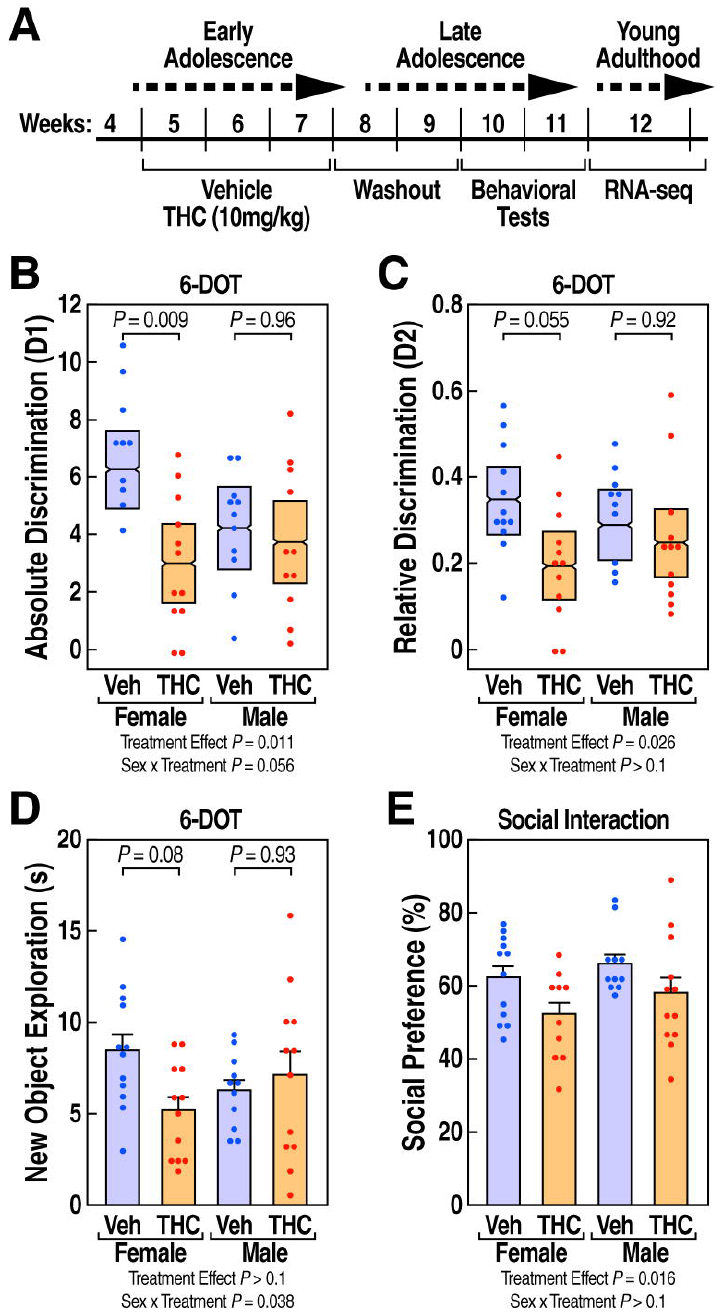
Adolescent exposure to THC reduced recognition memory and social interaction in a sex-specific manner. (A) Timeline of the study design. (B) absolute discrimination index D1 and (C) relative discrimination index D2 are shown as mean ± 95% confidence intervals showing decreased recognition memory in THC-treated groups compared to vehicle controls. (D) Exploration time(s) of the novel object is expressed as mean ± SEM showing a decrease only in female mice. (E) Social preference (%) is expressed as mean ± SEM and is reduced in both females and males. Main effects and interactions found using LMM analysis and posthoc comparisons p values using Tukey HSD test.

By using the six different objects test (6-DOT)^28,32^, we measured the effect of THC on object recognition memory. THC decreased the absolute (D1) and relative (D2) discrimination indexes by 36% (treatment effect F(1,41)=7.12, p=0.011) and 30% (treatment effect F(1,42)=5.3, p=0.026), respectively, compared to the vehicle control group (Fig 1B-C). There were no significant sex x treatment interactions for D1 (F(1,41)=3.86, p=0.056) or D2 (F(1,42)=1.89, p>0.1) indexes. However, exploratory posthoc analysis showed that, compared to the vehicle, THC decreased the D1 index by 44% (t=3.35, p=0.009) and the D2 by 39% (t=2.6, p=0.055) in females but not in males (p>0.1, Fig. 1B-C). Moreover, we observed a significant sex x treatment interaction (F(1,42)=4.6, p=0.038) in novel object exploration. A posthoc analysis revealed that THC-treated females, but not males, tended to reduce novel object exploration (t=2.44, p=0.08). The impact of THC was specific for the cognitive components of this assay, as THC had no detectable effect on the distance traveled during habituation (Fig. S1A) nor on the total time mice spent exploring the objects (all main effects and interactions p>0.1, Fig. S1B-C).

We also tested the effect of THC on social behaviors using the three-chamber interaction test. Adolescent exposure to THC significantly decreased social preference by 12% (F(1,41)=6.3, p=0.016) compared to the vehicle group (Fig. 1E). In contrast, sex had no detectable effect on treatment (sex x treatment F(1,41.5)=0.1, p>0.1).

Lastly, we examined anxiety-like behaviors with the elevated plus-maze. We did not observe significant differences in the willingness of mice to explore open environments (all main effects and interactions p>0.1, Fig. S 1D).

We excluded potential confounding effects on the exploratory activity by showing that THC treatment was not associated with changes in body weight at the time of behavioral testing (main effects and interactions p>0.1, Fig. S1E).

Overall, the behavioral analysis showed that exposure to THC during adolescence impaired memory and social interaction in late adolescence, with more robust memory deficits in female mice.

### Identification of DEGs associated with chronic adolescent exposure to THC

To gain insights into the neurobiological mechanisms underlying the behavioral alterations induced by THC, we profiled the transcriptome of PFC, DMS, NAc, Amy, and VTA from the vehicle and THC-treated mice (n=6/tissue/treatment/sex). Principal component analysis (PCA) validated the accuracy of brain dissection, showing that the transcriptomes of each brain region differed from one other (Fig. 2A). Furthermore, PCA plots within each brain region revealed substantial sex differences (Fig. S2A). In contrast, the effect of THC treatment was subtle (Fig. S2A). Next, we identified differentially expressed genes (DEGs) from each brain region and sex using different significance cutoffs (Fig. 2B, Table S1). We found the largest number of DEGs in the female Amy (n=743) and, in general, more DEGs in females in most brain regions except for NAc, which had the largest number of DEGs in males (n=77) at a false discovery rate (FDR)<0.1 and log fold change (logFC)>0.4 (Fig. 2B). Pathway analysis showed overlapping pathways altered by THC in female Amy and DMS, including opioid signaling and GPCR ligand binding (Fig. 2C). Moreover, DEGs in female Amy were also related to long-term potentiation, axon guidance, addiction, retrograde cannabinoid signaling, and translation (Fig. 2C). In contrast, DEGs in male NAc were involved in interferon signaling and ubiquitin-mediated proteolysis (Fig. 2C).

**Figure 2:**
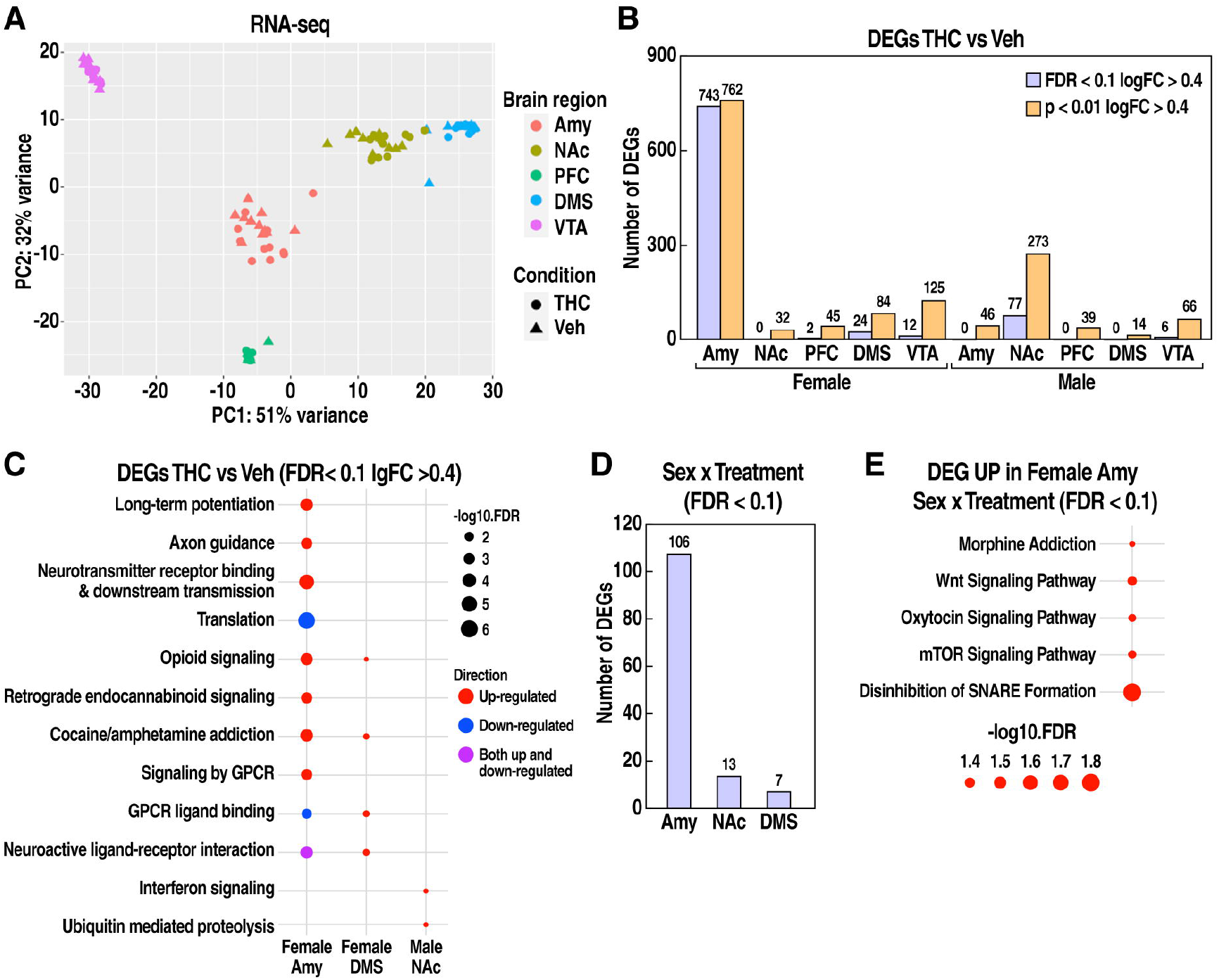
Adolescent THC exposure induced sex-specific transcriptional changes. (A) PCA visualization of male and female RNAseq samples across brain regions and treatment conditions. (B) Number of DEGs across brain regions and sexes using different significance cutoffs in analyses within each sex and brain region. (C) Pathway enrichment for DEGs in female Amy, DMS, and male NAc. Dot color depicts the direction of regulation and dot size illustrates the significance. (D) Number of DEGs for treatment by sex interaction across brain regions in analyses including both sexes for each brain region. (E) Pathway enrichment for genes upregulated by THC only in female Amy (sex x interaction). Dot size illustrates the significance.

To further explore the influence of sex on transcriptional responses to THC, we compared gene expression changes between males and females. At a statistical cutoff of FDR <0.1, this analysis yielded significant sex x treatment interactions for DEGs in Amy (n=106), DMS (n=7), and NAc (n=13), but not for PFC and VTA (Fig. 2D, Fig. S2B-D). Pathway analysis revealed that the genes upregulated by THC in the female Amy but not in males (Fig. S2B) were linked to presynaptic SNARE complex formation and several signal transduction pathways (e.g., mTOR, Wnt, oxytocin signaling (Fig. 2E). In contrast, other DEGs groups with sex x treatment interactions were not enriched in specific pathways (Fig. S2C-D).

In agreement with DEG analysis, threshold-free RRHO analysis showed minimal overlap in DEGs when we compared gene expression changes between most pairs of brain regions or between sexes (Suppl. Fig. 3).

These results indicate that females and males responded differently to THC in a brain region-specific manner.

### Identification of gene coexpression networks correlated with THC treatment and cognitive traits

Genes usually do not act alone but work as a network to achieve a biological function by interacting in a signal transduction or metabolic pathway ^33^. To better understand how THC impacts biological networks, we applied WGCNA^34^, a gene network modeling approach, to identify groups of genes (modules) highly coexpressed or coregulated in response to THC treatment within each brain region in each sex (Table S3). We then performed trait-module correlation analysis using cognitive behavioral traits measured in each mouse (Table S4). This analysis identified 27 modules across different brain regions significantly correlated (p<0.05) with THC treatment that we referred to as “THC-correlated modules” (Fig. 3A). While we did not observe any overlap between THC-correlated modules and those correlated with social preference, we identified five modules that simultaneously correlated with THC treatment and memory traits (Fig. 3B, S4A-B). Therefore, we will refer to these modules as “cognitive modules”. The cognitive modules included female DMS *saddlebrown* (Fig. 3C), female VTA *bisque4* (Fig. 3D) and *lightsteelblue1* (Fig. 3E), and male NAc *orange* (Fig. 3F) and *darkgrey* (Fig. S4C). Pathway enrichment analysis showed that DMS *saddlebrown* module was related to the metabolism of the endogenous cannabinoid anandamide and cognitive disorders, such as Alzheimer’s disease (Fig. 3C). The VTA *bisque4* module was enriched in interferon signaling and purinergic receptor genes (Fig. 3D), and the VTA *lightsteelblue1* module was involved in non-neuronal differentiation processes (Fig. 3E). In contrast, genes related to synaptic transmission were enriched in the male NAc orange module (Fig. 3F), but no pathway enrichment was identified for the male NAc *darkgrey* module. Notably, only female-specific modules showed a positive correlation with memory traits but a negative correlation with THC treatment (Fig. 3B). This observation suggests that the female cognitive modules regulate memory formation but are disrupted by THC treatment, reflecting the behavioral deficits observed in female mice. In contrast, the male NAc *darkgrey* showed positive correlations with both THC and memory traits but *orange* showed a negative correlation with both THC and memory traits (Fig. 3B), suggesting that the relationship between THC treatment and memory is more complex in males.

**Figure 3:**
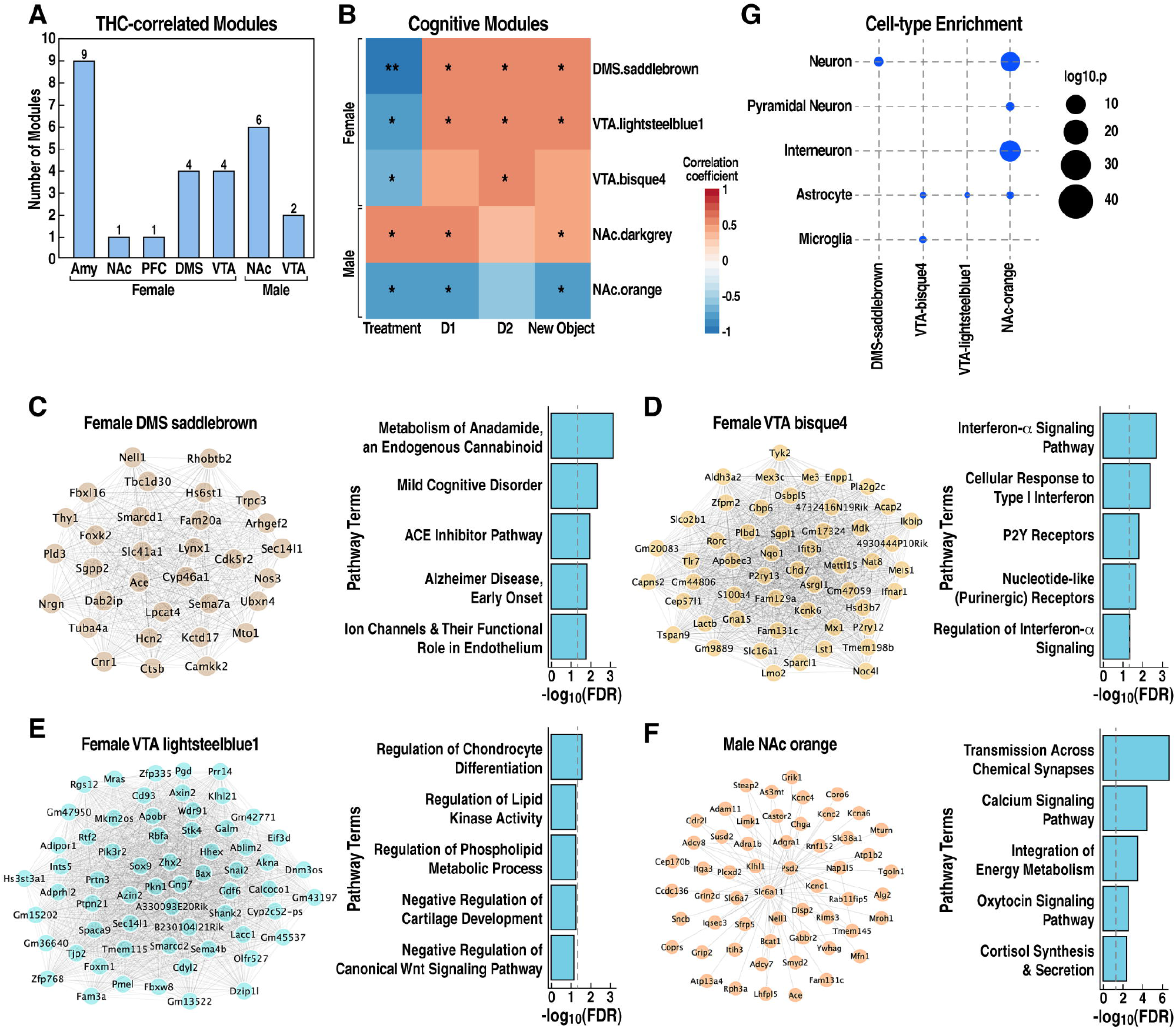
Characterization of cognitive modules correlated with THC treatment and mouse recognition memory. (A) Number of coexpression modules significantly correlated with THC treatment (p<0.05). (B) Heatmap of cognitive modules correlated with THC treatment and recognition memory. Color depicts the correlation coefficient with THC or memory traits. *p<0.05; **p<0.01; ***p<0.001. (C-F) Visualization of cognitive module networks and pathway annotations. The edges denote positive correlations between pairs of genes. Only the top 100 edges based on topological overlap weight were visualized due to the large size of the male NAc orange module. (G) Cell type marker gene enrichment of cognitive modules.

To identify the cell types that might contribute to the formation of cognitive modules, we performed cell-type marker enrichment analysis. In support of the pathway analysis, neuronal markers were enriched in the female DMS *saddlebrown* module and male NAc *orange* module, while markers of astrocytes and microglia were enriched in the female VTA *bisque4* and *lightsteelblue1* modules (Fig. 3G).

We found that THC-correlated modules and DEGs only partially overlapped (Fig. S5), indicating that network analysis captures additional information about THC transcriptional responses that go beyond changes in individual DEGs.

Overall, these results revealed that the effect of THC on memory is correlated with the regulation of sex- and brain region-specific gene coexpression modules.

### Cross-brain region module-module interactions affected by THC

During adolescence, dynamic changes in the eCBs coincide with the remodeling of circuit connectivity within and between brain regions, including corticolimbic structures^35–37^. To better understand the impact of THC on cross-brain gene coexpression networks in the female and male brains, we analyzed the correlations between modules within and between brain regions. This analysis identified numerous “THC-interconnected modules” that we define as those significantly correlated with THC-correlated modules in the same region or across brain regions with correlation coefficient |r|>0.5 and p<0.05 in females (Fig. 4A) and males (Fig. 4B). These modules likely reflect gene networks indirectly influenced by THC. Many THC-interconnected modules were also correlated with memory traits (colored nodes in Fig. 4A-B). As shown by the Sankey diagrams in Figure 4C-D, positive and negative correlations were relatively balanced across brain regions in both sexes. In females, higher levels of connectivity were observed between Amy-Amy modules, followed by Amy-VTA, VTA-VTA, DMS-DMS, and Amy-DMS (Fig. 4E). In males, higher levels of connectivity were observed between NAc-NAc, followed by NAc-VTA, NAc-PFC, VTA-NAc, and VTA-VTA (Fig. 4F).

**Figure 4:**
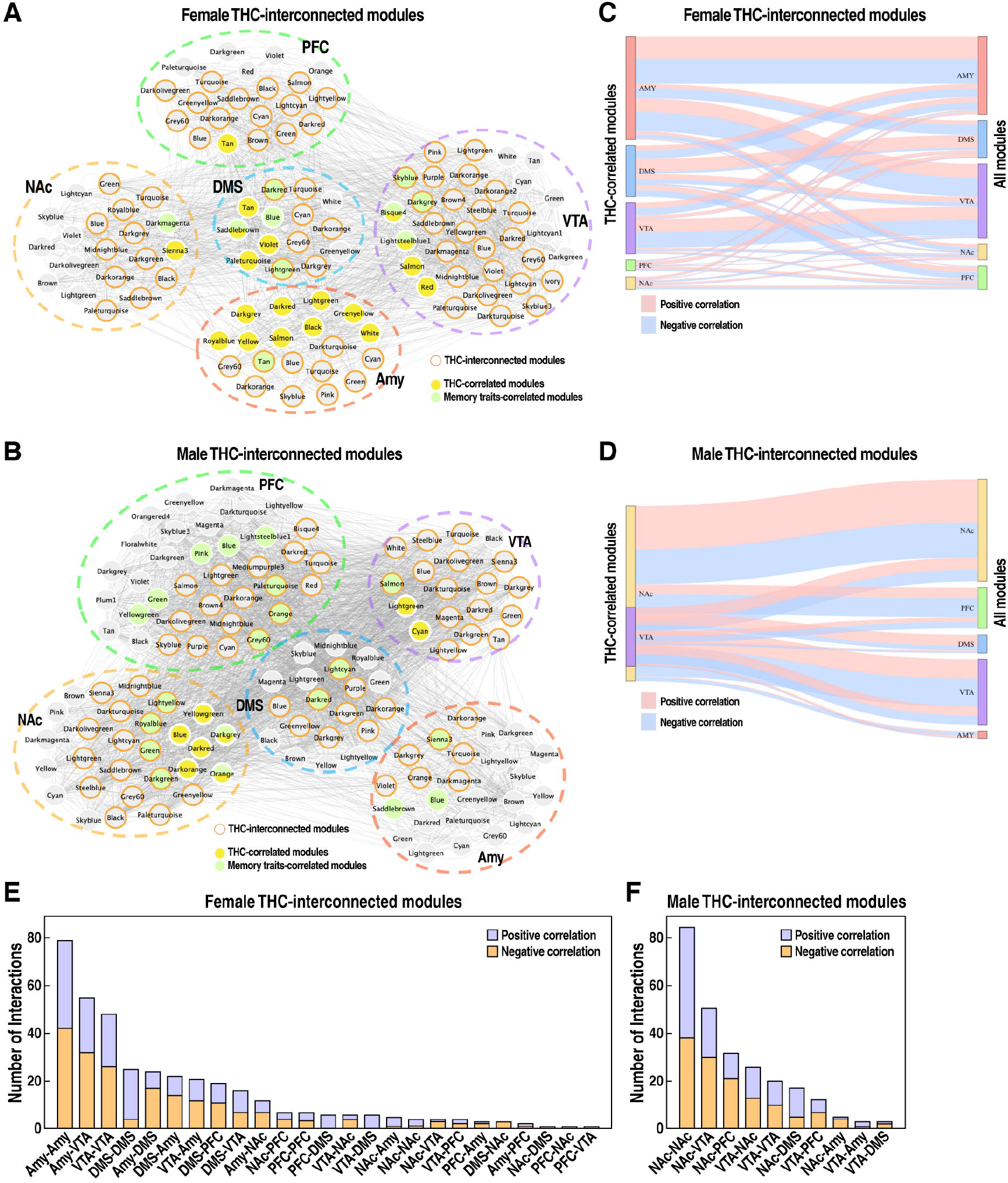
Construction of THC-interconnected module map reveals potential intra- or inter-region molecular circuitries disrupted by THC. (A-B) Visualization of female (A) and male (B) THC-interconnected modules, which are correlated with THC-correlated modules, with correlation coefficient |r|>0.5 and p<0.05. Nodes with filled colors denote modules correlated with THC or memory traits. Nodes with orange borderline depict THC-interconnected modules. (C-D) Sankey plots of female (C) and male (D) THC-correlated module interactions. Link colors denote the direction of correlation, with red indicating positive correlation and blue denoting negative correlation. (E-F) Number of intermodular interactions in THC-interconnected modules in females (E) and males (F). The color indicates the direction of correlation, with red indicating positive correlation and blue denoting negative correlation.

These findings suggest that adolescent exposure to THC leads to changes in molecular circuitries across different brain regions in a sex-specific manner.

### Associations between coexpression modules altered by THC and human cannabis use disorders

Recent GWAS have started to identify genetic variants associated with CUD^38^. To gain further insights into the genes and pathways associated with CUD, we applied the Mergeomic pipeline^31,39^ to integrate the human CUD GWAS signals with THC-correlated gene coexpression networks for each brain region and sex in mice (Fig. 5A). We defined “CUD-associated modules” as those enriched in CUD-associated genes informed by human GWAS (Table S6). There was no overlap between CUD-associated modules and THC-correlated modules in females or minimal overlap (11%, 2 modules) in males (Fig. 5B). In contrast, the overlap increased to 81.8% (9 modules) in females and 55.7% (11 modules) in males when we included THC-interconnected modules in the analysis (Fig. 5C). These results suggest that CUD-associated modules are likely indirectly affected by THC.

**Figure 5:**
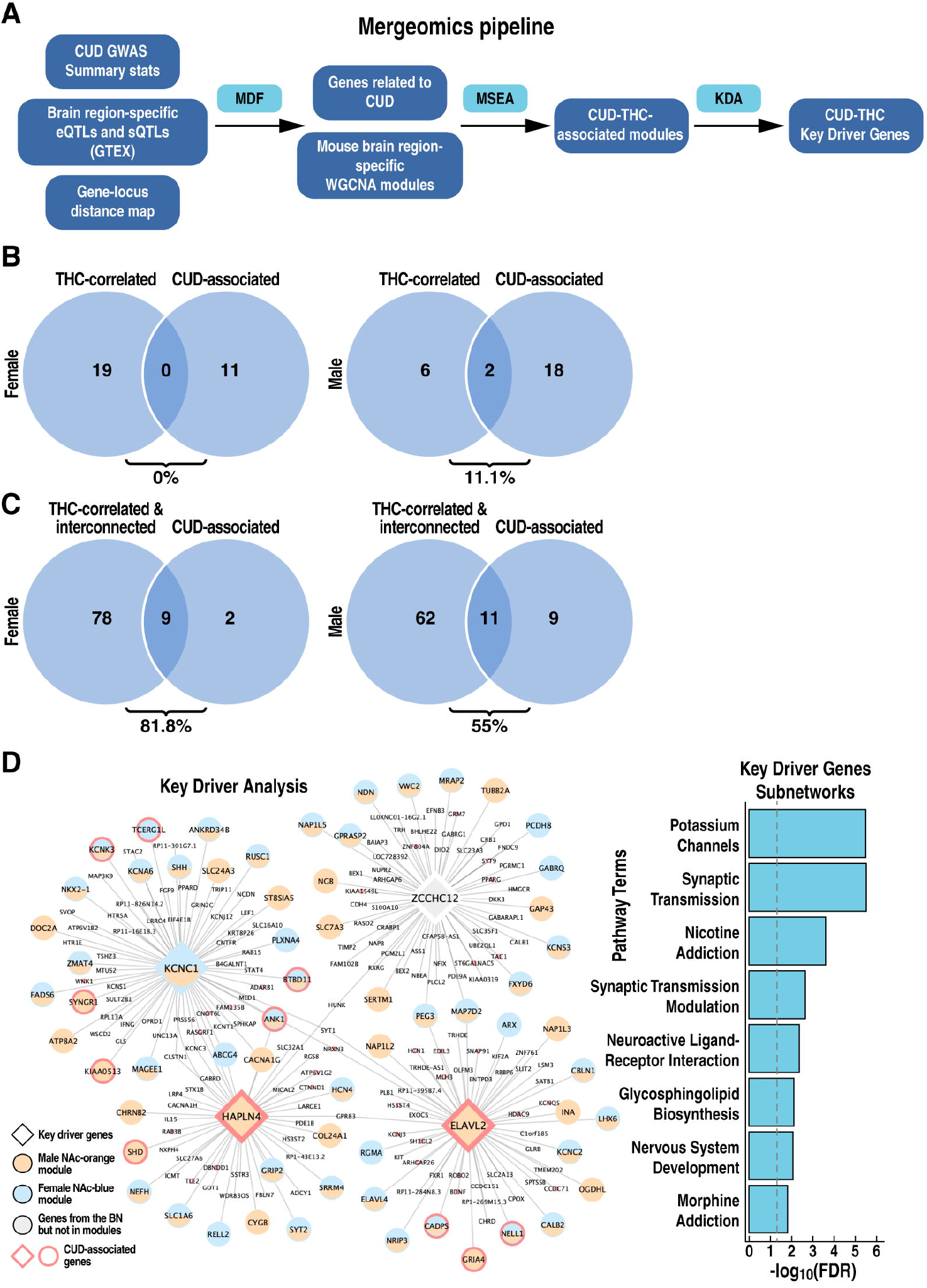
Association of THC-related modules with human CUD. (A) Schematic of Mergeomics pipeline. MDF: marker dependency filtering; MSEA: marker set enrichment analysis; KDA: key driver analysis. (B) Overlap (%) between CUD-associated modules and THC-correlated modules (C) or between CUD-associated modules and both THC-correlated and interconnected modules (D). The overlap (%) is calculated as the number of overlapping modules divided by the total number of CUD-associated modules. (D) Visualization of Bayesian network shared by CUD-associated modules in female and male NAc. Key driver genes are represented by large size diamond nodes. Orange, blue, and grey nodes denote genes (male NAc orange module, genes in female NAc blue module, and genes in the BN but not in the two aforementioned modules, respectively. CUD-associated genes identified by Mergeomics using loci with p<0.001 from the Johnson et al.^11^ CUD GWAS are labeled with red borderline. Bar plot depicts pathway enrichment of the genes in the CUD subnetwork. BN, Bayesian network

To predict potential key regulators of CUD-associated networks, we performed key driver (KD) analysis using tissue-specific Bayesian networks that infer causal relationships between genes and that were constructed using independent human and mouse data^39^. We identified top KD regulators of gene coexpression networks associated with THC and CUD (overlapping modules in Fig. 5C) in males and females (Table S7). While most brain regions showed sex-specific KD genes, the NAc showed an overlap of four KD genes between males and females (*Hapln4, Kcnc1, Elavl2, Zcchc12*, Fig. 5D), suggesting a common regulatory mechanism linked to CUD. Among these genes, two encode for membrane proteins involved in the modulation of synaptic plasticity. KNCN1 is a voltage-gated potassium channel ^40^, and HAPLN4 is a component of the perineuronal net^41^. The other two KDs encode proteins involved in transcriptional regulation. ELAVL2 is an RNA-binding protein involved in splicing in neuronal development^42^, and ZCCHC12 is a neuronal transcriptional coactivator^43^. Of note, two KD genes, *Hapln4* and *Elavl2*, were also identified as CUD-associated genes. The pathways analysis of the four KD-associated subnetworks shared between males and females revealed that they regulate genes implicated in addiction processes, neurotransmission, brain development, and lipid metabolism (Fig. 5D).

This integrative genomic analysis identified a connection between genes and pathways altered by THC and associated with CUD vulnerability.

## Discussion

Our work provides the first comprehensive, tissue- and sex-specific view of molecular processes perturbed by adolescent THC treatment in mice and linked to CUD in humans (Fig. 6). We identified gene coexpression networks disrupted by THC in specific brain regions and correlated to memory deficits induced by THC in a sex-specific manner. In addition, we identified key regulators that orchestrate brain region-ecific transcriptional subnetworks linked to adolescent exposure to THC and CUD vulnerability.

**Figure 6:**
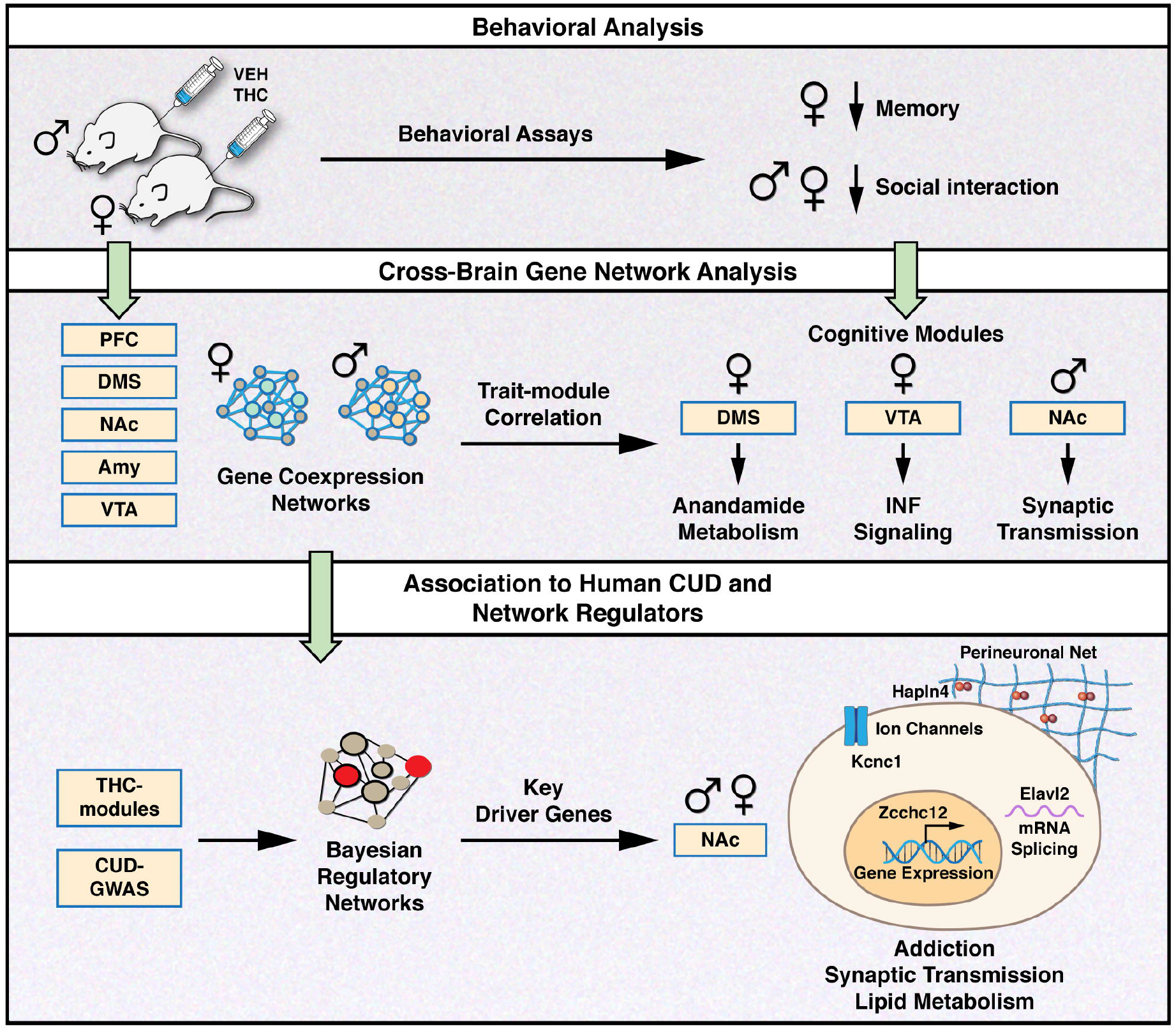
Integrative genomic analysis identified sex- and brain region-specific biological processes and regulators affected by THC and linked to CUD. (Top panel) Behavioral analysis showed that adolescent exposure to THC leads to cognitive impairments, with stronger memory deficits in female mice. (Middle panel) Gene coexpression analysis identified sex- and brain region-specific cognitive modules correlated with THC treatment and memory impairments. (Bottom panel) The integrative genomic analysis identified potential causal regulators of gene networks associated with THC treatment and CUD vulnerability.

In line with previous reports in rodent models^28,44–50^, our behavioral analysis demonstrated that adolescent exposure to THC in mice led to long-term impairments in object recognition memory and social interaction, but not in anxiety-like behaviors. Our study also showed sex differences in the effects of THC on recognition memory, which was impaired more in females compared to male mice. Although previous preclinical studies have not examined the influence of sex on the effects of THC on memory, female rats have been reported to be more susceptible than males to the effects of THC on locomotor activity, nociception, and reward processes^51–55^. In addition, clinical studies showed that females are more sensitive to the harmful effects of THC on spatial memory^56–60^.

In agreement with the sexual dimorphism observed for the behavioral effects of THC, we also reported extensive sex differences in gene expression patterns in response to THC. First, female mice showed a larger number of DEGs across different brain regions compared to males. Secondly, when we conducted a statistical analysis that explicitly tested for sex differences in DEGs, we found significant sex x treatment interaction in Amy, NAc, and DMS, suggesting that females and males respond differently to THC. These results are new, as prior research on the transcriptional effects of adolescent exposure to THC has focused only on male rats^26,27^. Miller et al reported that adolescent exposure to THC in male rats was associated with gene expression changes related to cytoskeleton and chromatin regulation in the PFC^26^. In contrast, we did not identify any DEGs or THC-correlated modules in the male PFC in mice. Between-species differences our different statistical thresholds in DEG analysis may explain the discrepancy.

Which brain region drives distinct behavioral abnormalities induced by adolescent exposure to THC is not entirely known. Our work, for the first time, simultaneously investigated five brain regions. Our results indicated an extensive brain region specificity in the genes and networks altered by THC. Amy and NAc may be sites of particular importance as they were associated with the largest number of DEGs in females and males, respectively. In line with these findings, brain morphological studies of human cannabis users have shown that marijuana use may be associated with disrupting the neural organization of the Amy and NAc^61^. In particular, previous studies have documented that female teenagers who use marijuana are more susceptible than males to structural abnormalities of the Amy, which were correlated with worse internalizing symptoms^62^. Consistent with these observations, animal studies also reported perturbation of synaptic transmission in the Amy and NAc following administration of exogenous cannabinoids^63,64^.

Moreover, our gene co-expression analysis also identified specific brain regions linked to memory traits in mice. Specifically, the disruption of female cognitive modules in DMS (*saddlebrown*, enriched for eCB related pathways) and VTA (*bisque4* and *lightsteelblue1*, enriched for immune and non-neuronal differentiation pathways) was correlated with the deficits in recognition memory observed in female mice. In agreement with the pathway annotation of these modules, it is well known that the endocannabinoid system plays an essential role in learning and memory processes, with the engagement of the dorsal striatum specifically in encoding habit-related memories^65–67^. Moreover, repetitive exposure to synthetic cannabinoids led to inflammatory phenotypes, including astrogliosis in the VTA^68^.

Our multiple brain region studies also allowed us to uniquely infer network connections within and between brain regions. The THC-interconnected modules are likely indirectly influenced by THC, as inferred from the module-module interaction network. We speculate that the inter-region interaction network could predict how THC directly affects one brain region that then cascades down to other brain regions. For example, our analysis suggests that adolescent exposure to THC alters neural circuits that connect Amy with VTA and DMS in females and neural circuits that connect NAc with VTA and PFC in males. Future experiments perturbing the THC modules using animal models will be necessary to validate these predictions.

Cannabis use disorder has a strong genetic component and is influenced by other environmental factors, including social and developmental vulnerability. For example, early initiation age in adolescence and a high frequency of cannabis use increase the risk of CUD^10^. Our integrative genomic analysis identified CUD-THC subnetworks and potential causal regulators, including four KD genes (*Hapln4, Kcnc1, Elavl2, Zcchc12*) shared between male and female NAc and implicated in addiction processes, synaptic transmission, brain development, and lipid metabolism. However, the involvement of these genes in CUD has not been explored before, and follow-up studies will be needed to confirm the role of the key driver genes in mediating THC effects on CUD in vivo.

Our results should be considered in light of certain limitations. First, we focus on correlating transcriptomic changes that occur in late adolescence with behaviors measured at the same time. The advantage of this approach is that it can capture gene expression changes associated with a history of early adolescent exposure to THC. However, it cannot directly assess the transcriptional and behavioral changes occurring while the drug is onboard. The second limitation of our study is that we focus on cognitive behaviors, such as recognition memory and social interaction. Still, other behaviors are likely to be influenced by THC, including addiction-like phenotypes. However, it is important to note that the addictive properties of THC are not well modeled in mice. Thirdly, our study is limited to 5 brain regions and can miss other additional gene regulatory networks. Thus, it would be important to expand this study to other brain regions, such as the hippocampus, given its critical role in recognition memory. Lastly, we cannot exclude that mechanisms other than gene expression changes contribute to sex-specific THC-related behaviors. For example, sex-specific hormonal changes^69^ or pharmacokinetic factors^70^ during adolescence and the differential density of cannabinoid receptors in the female and male brain^71^ could contribute to the sex differences observed in our study. Our findings open numerous new hypotheses that warrant future experimental validation.

In summary, our study is the first to integrate gene expression profiles, GWAS, and network modeling to reveal comprehensive sex- and brain region-specific view of biological processes and regulators influenced by cannabis use and linked to CUD vulnerability.

## Acknowledgments

This work was supported by the National Institute on Drug Use, USA [DP1DA042232, U01DA050239 to FT]. XY is supported by the National Center for Advancing Translational Sciences UCLA CTSI Grant UL1TR001881.

This publication includes data generated at the UC San Diego IGM Genomics Center utilizing an Illumina NovaSeq 6000 that was purchased with funding from a National Institutes of Health SIG grant (S10OD026929)

We would like to acknowledge H. Taylor and J. Hightower for technical assistance.

## Conflict of Interest

The authors have no conflicts of interest to disclose.

Supplementary information is available at MP’s website

---

**Supplementary Figure 1:**
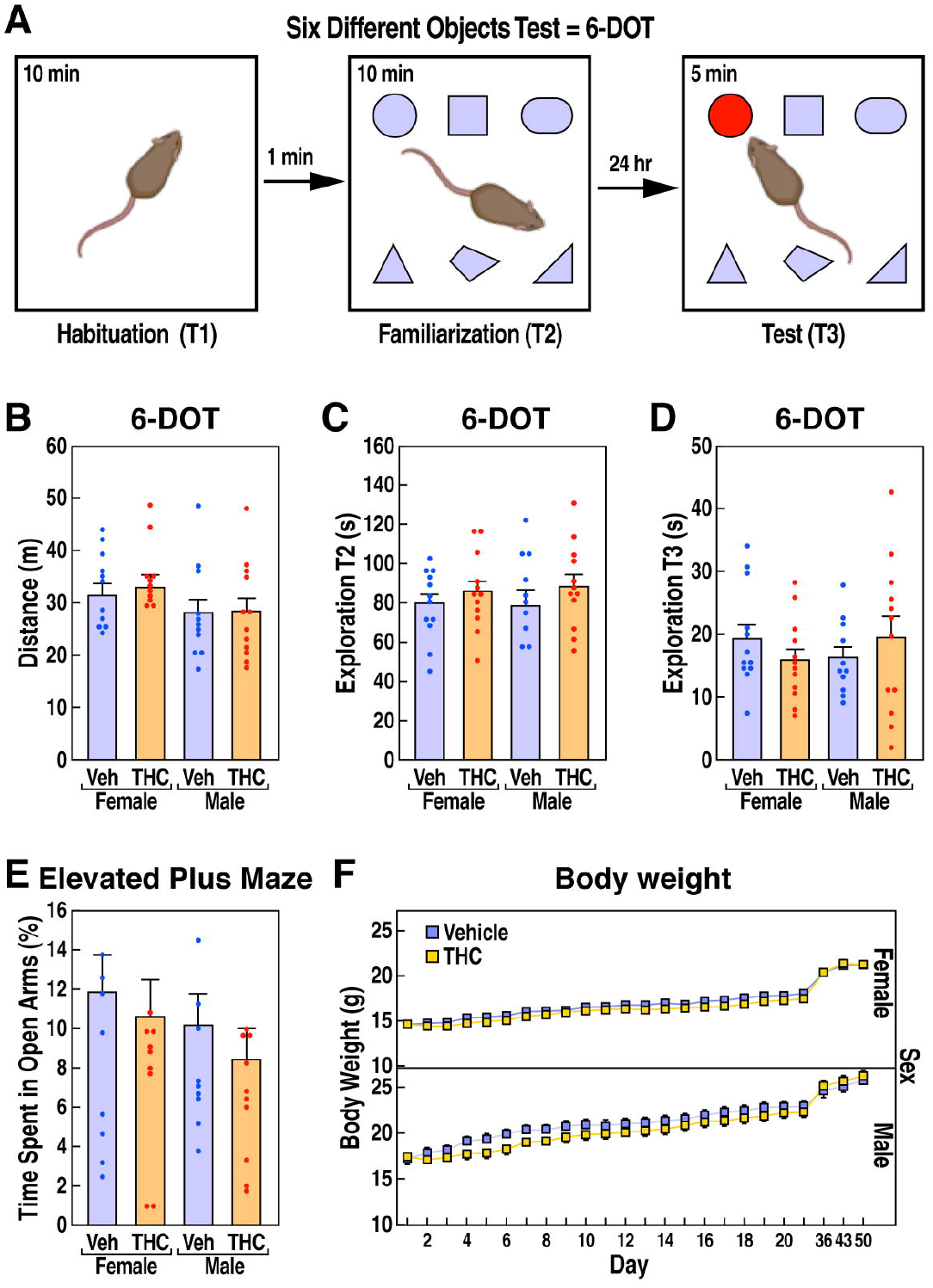
Behavioral characterization of female and male mice following adolescent exposure to THC. **(A)** Locomotor activity is expressed as mean distance traveled (m) ± SEM. Exploration (s) in familiarization phase T2 **(B)** and test phase T3 **(C)** are expressed as mean ± SEM. **(D)** Bodyweight (g) is expressed as mean ± SEM for the THC administration course (day 1 to 21) and day 36, 43, and 60 when the three behavioral assays started. Main effects and interactions found using LMM analysis, and posthoc comparisons p values using Tukey HSD test.

**Supplementary Figure 2:**
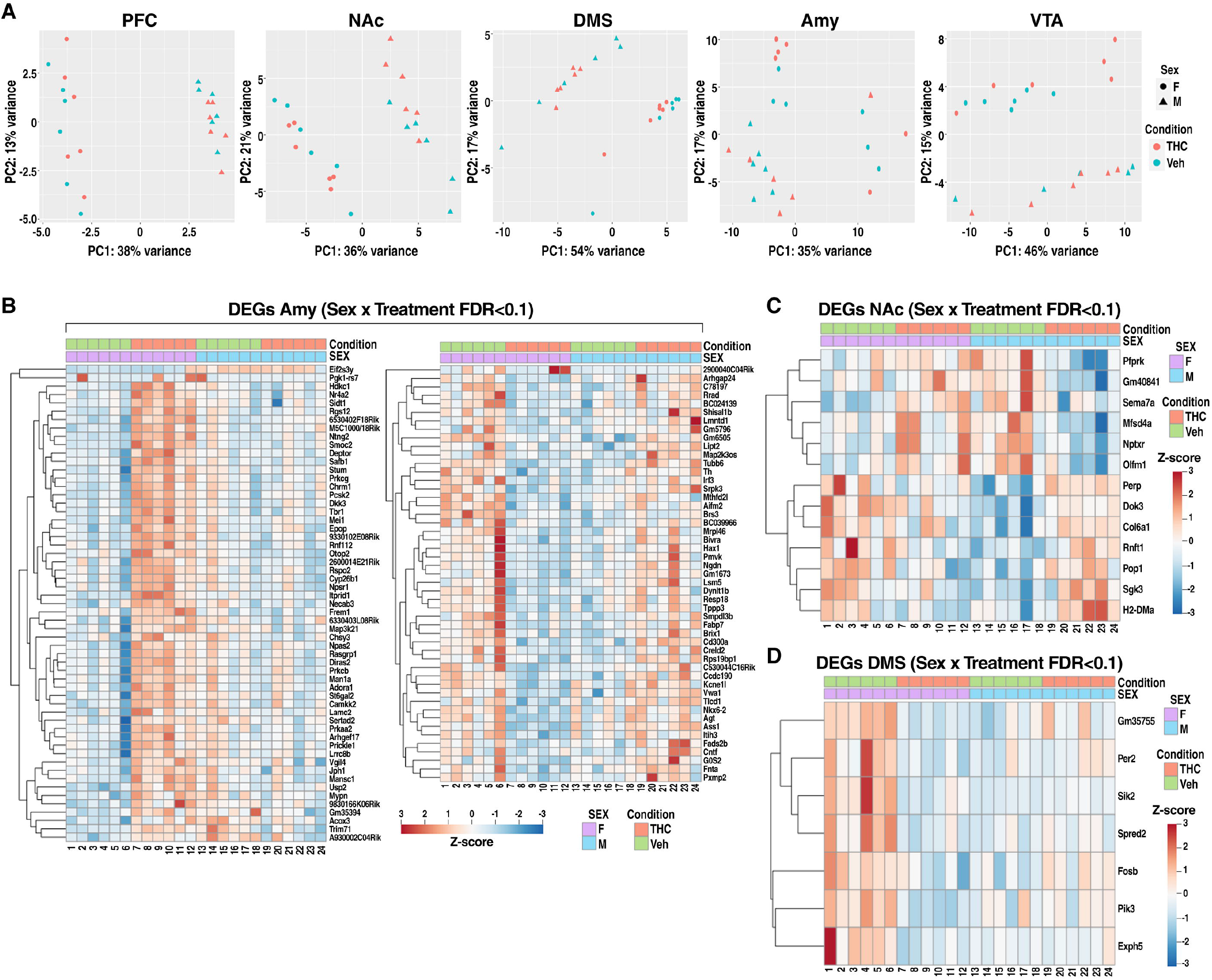
Adolescent THC administration induced sex-specific transcriptional changes. (A) PCA visualization of all samples in each brain region. Samples are colored with sex and treatment conditions. (B) Heatmaps of treatment by sex interaction DEGs in Amy. The left panel depicts DEGs upregulated in females with THC treatment, while the right panel depicts DEGs upregulated in males with THC treatment. (C) Heatmaps of treatment by sex interaction DEGs in NAc. (D) Heatmaps of treatment by sex interaction DEGs in DMS. UP, upregulated; DN, downregulated.

**Supplementary Figure 3.**
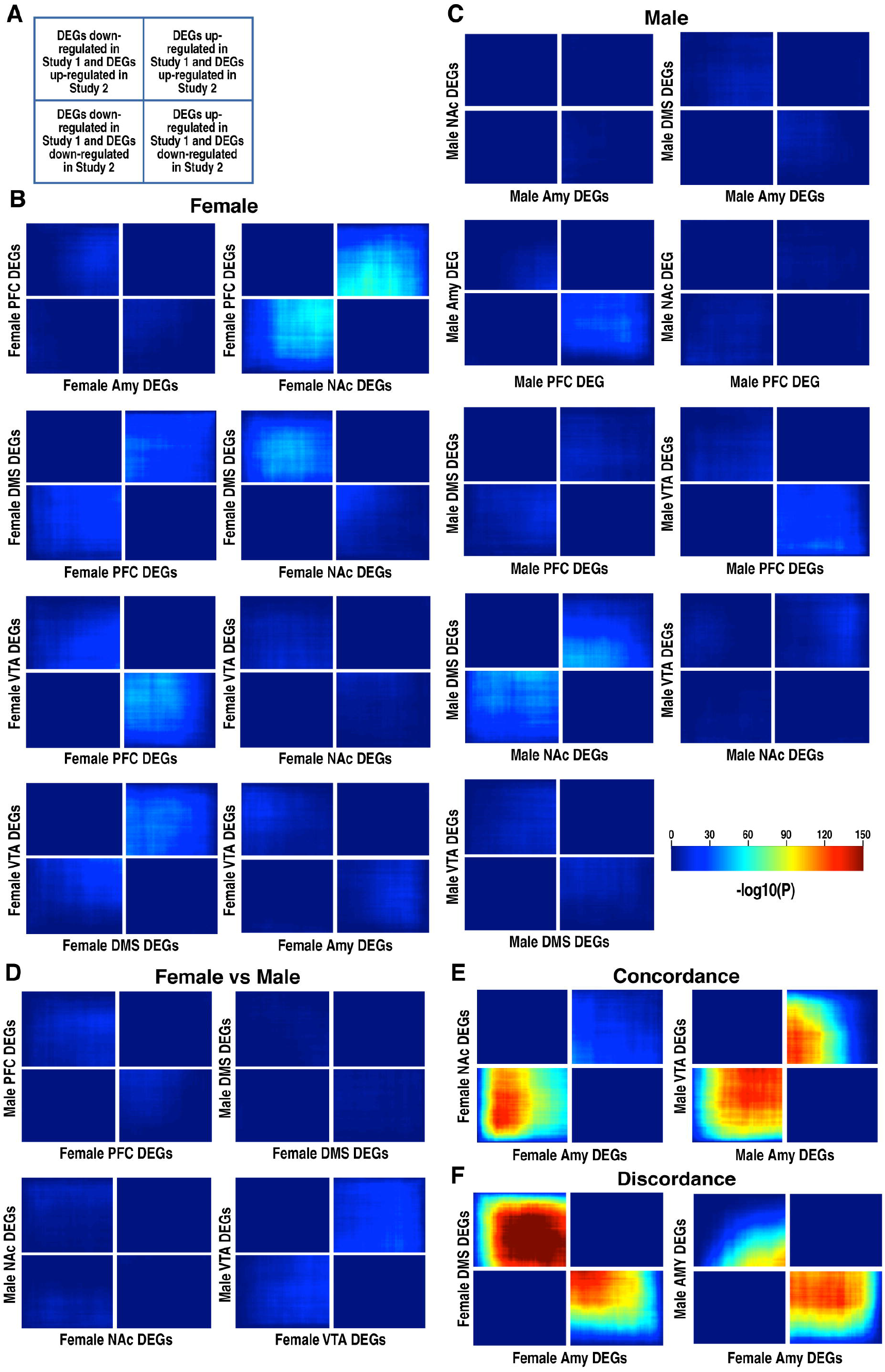
Minimal transcriptional overlap was detected in most comparison pairs across sex and brain regions based on rank-rank hypergeometric overlap (RRHO) analysis. (A) Schematic of RRHO heatmap interpretation. (B-D) RRHO heatmaps show minimal overlap in transcription between pairs of brain regions within females (B), males (C), and between sexes (D). (E-F) RRHO heatmaps show concordance (E) and discordance (F) in transcription between pairs of brain regions. We define concordant as positive correlation and discordant as negative correlation in DEG overlap patterns.

**Supplementary Figure 4:**
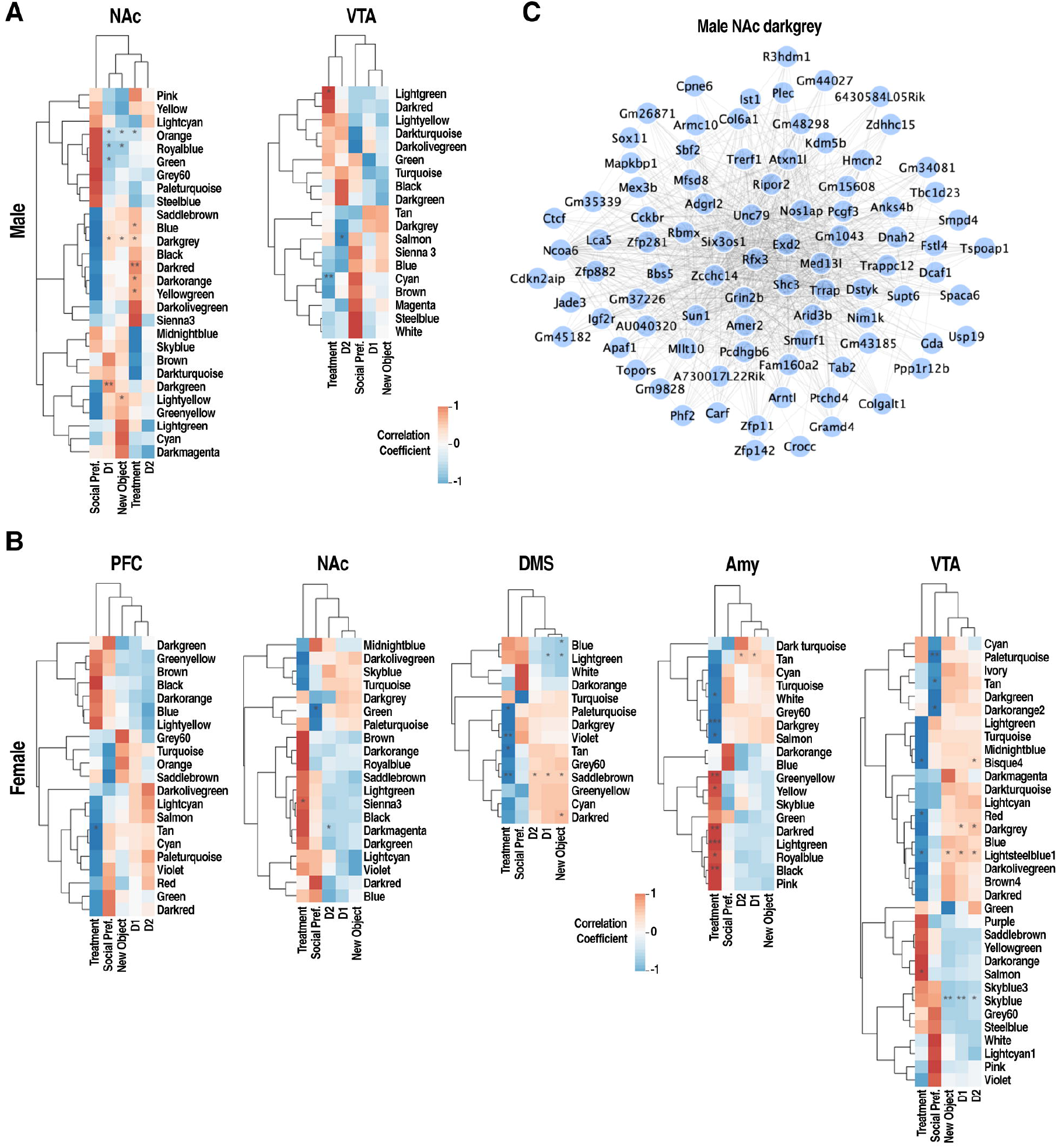
DEG enrichment in WGCNA coexpression modules was detected in a few brain regions. (A-B) Heatmaps of DEG enrichment in WGCNA coexpression modules in individual brain regions. Select modules from female Amy, DMS, PFC, VTA and male VTA, and NAc showed significant DEG enrichment. Female Amy had the largest number of modules that showed DEG enrichment. Color depicts -log10 (P value) of the enrichment by Fisher’s exact test. *p<0.05; **p<0.01; ***p<0.001. DEG.all, DEG.up, and DEG.down denotes all, up-regulated, and down-regulated DEGs of a certain brain region, respectively.

**Supplementary Figure 5:**
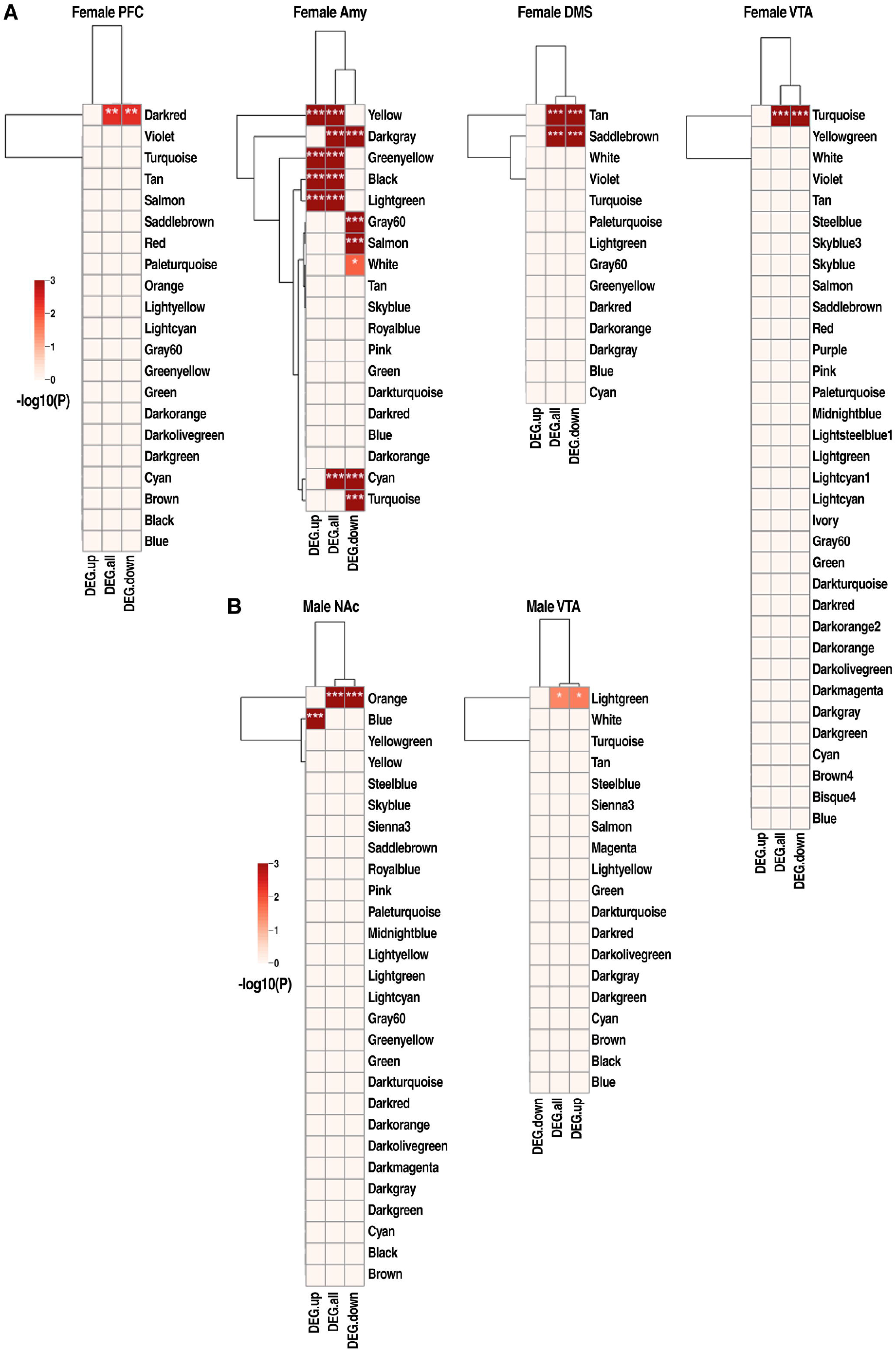
Correlation of WGCNA coexpression modules with chronic THC administration and mouse cognitive behavioral traits. (A-B) Heatmaps of module correlation with THC treatment and cognitive traits in males (A) and females (B), respectively. Only brain regions with significant correlation are shown. Color depicts correlation coefficient relative to THC or memory traits. *p<0.05; **p<0.01; ***p<0.001. (C) Visualization of male NAc darkgrey module. The edges denote positive correlations between pairs of genes. No pathway annotation under the significance of FDR < 0.05 was found.

