## Supplemental Methods for "Chronic adolescent exposure to cannabis in mice leads to sex-biased changes in gene expression networks across brain regions"

**Supplementary Methods and Materials**

**Animals**

C57BL/6NCrl male (n=24) and female (n=24) mice were purchased from Charles River (Strain Code: 027). Mice were housed (3-4 per cage) under a 12h light/12h dark cycle and provided with food and water ad libitum. All experimental procedures were approved by the institutional animal care and use committee at the University of California, San Diego.

**Drug treatment protocol and experimental design**

THC was provided by the U.S. National Institute on Drug Abuse and was dissolved in a vehicle solution consisting of ethanol, tween, and 0.9% saline (1:1:18) on the day of administration. As shown in Fig. 1A, our experimental design covers different developmental periods, including early adolescence (5-7 weeks), late adolescence (8-11 weeks), up to young adulthood (~ 12 weeks) in rodents(1–4). Specifically, we administered vehicle or THC (10 mg/kg) daily to mice by intraperitoneal injections in the early adolescent period, from postnatal day (PND) 28 to PND 48 (week 5-7), as also

previously reported(5,6). In our previous study, we used the same drug treatment

protocol and showed that after daily injections of 10mg/kg THC for three weeks starting at PND 48 (5 weeks old), the plasma concentration of THC was 5-6 ng/ml one day after the last injection. Although less is known about plasma THC concentration in adolescent users, these results are consistent with previous studies in chronic frequent adult cannabis smokers which reported THC in plasma with a median ≥5.0 ng/ml up to 3 days of abstinence (7–9). Although there is a large variability in the amount of THC in cannabis used by humans, the dosage used in the current study likely represents heavy use(7–9).

To study the behavioral effects of adolescent exposure to THC, we tested 2 groups of mice for behavioral assays (n = 24 mice/group) two weeks after the last THC or vehicle injection, starting at PND 63 (week 10). Behavioral assays were conducted on separate days and all behavioral tests were performed once on each mouse. To prevent bias due to olfactory cues, the behavioral apparatus was cleaned in between mice. Bodyweight (g) was measured through the course of the drug administration protocol and when each behavioral assay began. At the end of the behavioral protocol (week 12), brains from one group (n = 24 mice) were collected in dry ice and stored at -80^o^C before dissections and molecular analysis.

**Elevated Plus-Maze (EPM) Test**

The EPM was performed as described in Iemolo et al.(10). Briefly, mice were placed individually onto the center of the maze for a 5-min period. The percent of total arm time [i.e.,100 × open arm/(open arm + closed arm)] was measured and used as an index of anxiety-related behavior(11).

**Six different objects test (6-DOT)**

We used the 6-DOT (Fig. S1A) to study recognition memory in mice, as previously described(6) with minor modifications. Mice were left free to explore an empty arena during a 10 min habituation trial (T1). Afterward, mice were exposed to six different objects for 10 min (familiarization trial, T2). After 24 hours, mice were exposed for 5 min to 5 familiar objects and a novel object (test trial, T3). The exploration time for each trial was measured with Anymaze (Ugo Basile, Varese, Italy). Absolute discrimination index D1 = (time spent exploring novel object – average time spent exploring familiar objects), and a relative discrimination index D2 = [(time spent exploring novel object – average time spent exploring familiar objects) / Total time spent exploring novel + familiar objects]. The D2 index also takes into account differences in exploration time between mice(12). The discrimination indexes were used as measures of recognition memory. A longer time spent exploring the novel object is also interpreted as a sign of memory reconsolidation(13) or novelty-seeking behavior(14). During the habituation trial, total distance traveled (m) was measured to assess basal locomotor activity.

**Three-chamber social approach test**

The three-chamber social approach test was performed as previously described(6). The target mouse was first placed in the center chamber and allowed to explore the apparatus for 15 min. After introducing the novel mouse and the novel object in the side chambers, the mouse was allowed to explore for 10 min. The placement of the novel mouse or novel object in the left or right chambers was systematically alternated in between trials. The time spent actively sniffing the novel mouse or the novel object was manually scored. A longer time exploring the novel mouse versus the novel object was considered an index of sociability. Social preference was calculated as [(Time sniffing novel mouse/(total time sniffing novel mouse and novel object))*100].

**Statistical analysis of behavioral measures.**

To evaluate how factors (treatment, sex) influenced mice behavior, we used linear mixed models (LMM) in JMP pro v. 16.0 (SAS Institute, Inc). LMMs allow both fixed and random effects, which include repeated measures(15). We modeled the categorical predictor variables for the overall effect of treatment and sex, and we included sex x treatment interaction to study sex-specific effects. We used mouse cohorts and individual subjects as random effects (when appropriate) to account for possible non-independence of the data. Tukey HSD post hoc pairwise comparisons identified differences among specific groups. A p < 0.05 was considered significant. Outliers were detected using the Huber M-estimation method(16) in JMP pro 16.0 and removed when appropriate (*n* = 1 in absolute discrimination index D1 and *n* = 1 in social preference).

**Brain punching**

Brains were sliced on a cryostat, and 5 brain regions were collected using 1-2mm punches, starting at the following bregma coordinates: +2.68 (PFC), 1.78 (NAc), 1.10 (DMS), -1.34 (Amy), and -2.92 (VTA), for a total of 120 samples.

**RNA extraction**

Brain punches (n = 120) were homogenized in Trizol Reagent (Invitrogen, Cat, num. 15596018) and Zirconium Beads RNase Free (Next Advance, Cat. num. ZrOB05-RNA 0.5mm) using the Bullet Blender Blue (Next Advance, Model. num. BBX24B) at speed 6 for 1 min. RNA was then purified using the Direct-Zol kit (Zymo, Cat. num. R2052). RNA quality was assessed using the Agilent 2200 TapeStation system and samples with RIN above 7.5 were used for library preparations. One sample for a VTA vehicle-treated male mouse with low RIN was excluded.

**RNA-seq library preparations**

An average of 200ng of RNA was used for each sample. The NEBNext Poly(A) mRNA Magnetic Isolation Module (New England BioLabs, Cat. No. E7490L) and the NEBNext Ultra II Directional RNA Library Prep Kit (New England BioLabs, Cat. No. E7760S) were used following the manufacturer’s instructions. A total of 119 libraries were sequenced on the Illumina NovaSeq 6000 system (2 x 100 bp reads), which generated an average of 27.7 million reads/sample.

**RNA-seq raw data processing and identification of differentially expressed genes (DEGs)**

Raw sequencing reads were cleaned to remove adapter sequences and low-quality sequences using cutadapt v2.9(17) with the paired-end trimming mode and the minimum 18-base setting. The cleaned reads were aligned to the mouse genome (*mm10*) using STAR v2.7.3a(18) following ENCODE parameters (<https://github.com/ENCODE-DCC/long-rna-seq-pipeline/blob/master/DAC/STAR_RSEM.sh>), which generated an average of 80% alignment rate. The aligned reads were quantified at the gene level using RSEM v1.3.1(19). Principal component analysis (PCA), Cook’s distance, and hierarchical clustering were used to detect potential sample outliers. One sample from male VTA was detected and removed from further analysis. The quantified gene-level counts were analyzed with DESeq2 v1.28.1(20) for differential gene expression analysis by comparing the THC- versus vehicle-treated groups for each brain region and sex. Significant DEGs were identified using FDR <10% and absolute log (fold change) > 0.4 as the cutoff. To identify sex x THC treatment interaction DEGs, we performed DEG analysis using the following formula for each brain region: gene expression ~ sex+treatment+sex:treatment. Significant sex:treatment interaction DEGs were identified using FDR < 10% as the cutoff.

**Rank Rank Hypergeometric Overlap (RRHO) Analysis**

To assess consistency or disagreement in the gene expression changes induced by THC between brain regions within each sex, or between males and females for each brain region, we performed RRHO analysis using the RRHO2 R package(21,22). RRHO analysis detects the overlap of transcriptional profile change in the same or opposite directions between two datasets using a threshold-free approach. Full differential expression signatures were used to generate a ranking score based on the following formula: score = - log10(P-value) * sign(log2FoldChange). Alignment of DEG signature between each pair of brain regions was examined in males and females, respectively. A comparison of DEG signature alignment between sexes for each brain region was also conducted. RRHO heatmaps were generated for each comparison using the RRHO2_heatmap function in the RRHO2 R package.

**Weighted gene coexpression network analysis (WGCNA)**

RNA-seq data was normalized with DEseq2 median of ratios normalization and then log-transformed, and genes with a standard deviation larger than 0 were chosen for WGCNA analysis. The WGCNA R package was used to generate ten signed coexpression networks for the PFC, NAc, DMS, Amy, and VTA for males and females separately (23). An adjacency matrix was generated to capture pair-wise correlation relationships in terms of biweight midcorrelation between genes, using dataset-specific soft power for the best fit of a scale-free topology. A topological overlap measure (TOM) matrix was then calculated based on the adjacency matrix to measure relative interconnectedness between each pair of genes. Next, a gene dendrogram was built using the flashClust function based on hierarchical clustering. Branches of the dendrogram correspond to clusters of positively correlated genes, which are termed modules. The cutreeDynamic function based on a branch-cutting algorithm were used to identify modules using the following parameters: method=hybrid, minModuleSize = 30, pamStage = F, pamRespectsDendro = T. DeepSplit and cutHeight value were chosen on a per-network basis to optimize module detection. After module detection, module eigengenes (MEs) were calculated as the first principle component of genes in a module to summarize each module’s expression profile. Modules with ME correlation higher than 0.85 were considered highly similar modules and merged using the mergeCloseModules function. To examine the similarity of each gene’s expression profile with its module’s ME, biweight midcorrelation to the ME was calculated for each gene as its module membership score. In each module, hub genes were defined as genes with an intramodular connectivity score among the top 10% in the module and a gene module membership score higher than 0.8. To identify module-trait associations, Pearson correlations between module MEs and treatment, social preference score, object discrimination indexes D1 and D2, and new object interaction time were calculated. Modules with significant correlation (p < 0.05) with one or more traits were used for weighted key driver analysis to identify key driver genes in a Bayesian network, as described below in Weighted Key Driver Analysis (wKDA).

**Pathway annotation of DEGs, WGCNA coexpression modules, and subnetworks**

Pathway annotation for the DEG sets and WGCNA modules from individual brain regions was performed using the enrichR R package(24). General databases including KEGG, Reactome, GO Biological Process, BioCarta and WikiPathways from enrichR’s database library were used for DEG pathway, WGCNA module, and subnetwork annotations. Pathway enrichment was calculated based on the deviation from the expected rank for terms, and P-values were adjusted for multiple testing corrections using Benjamini-Hochberg correction. Adjusted P values of 0.05 were considered significant.

**DEG and cell type marker enrichment in WGCNA modules**

To explore the potential cell type source for the cognitive modules, we used cell type markers curated from Werling and coauthors(25) to perform enrichment analysis, including pyramidal neurons, interneurons, oligodendrocytes, microglia, activated microglia, astrocytes, mural cells, endothelial cells, and ependymal cells. Between each cell type marker set and each WGCNA module, a fold enrichment score and enrichment p-values were calculated using the phyper R function. We assessed the overlap between the WGCNA modules and DEGs. DEGs with an FDR value lower than 0.1 and the absolute value of logFC equal to or higher than 0.4 were extracted for overlap analysis. DEGs were further split into upregulated and downregulated DEGs based on whether the corresponding logFC was positive or negative. Each DEG set was assessed for overlap with WGCNA modules using the method described for cell-type marker overlap above.

**Weighted Key Driver Analysis (wKDA)**

wKDA from the Mergeomics R package was used to identify key drivers of selected modules or DEGs based on a gene interaction network(26). We used a brain-specific Bayesian network capturing causal gene-gene regulatory relationships constructed using the RimbaNet package(27) and numerous human and mouse brain genetic and transcriptomic datasets(28). Modules with a significant correlation with one or more traits as well as DEGs for each brain region were extracted and used as input for wKDA. In wKDA, hub genes were first identified based on the input network’s topology. Subsequently, the input module genes or DEGs were overlaid onto the network. The hub genes whose downstream network neighbors were enriched with the input module genes or DEGs were highlighted as key drivers for the trait-related genes or modules. The parameters used were as follow: edgefactor=1 (considering edge weights), depth=1 (searching depth), direction=0 (not considering the directions of the interactions), nperm=10000 (permutation numbers), ndrivers=10 (up to 10 key drivers from each input gene list). Networks of prioritized key drivers were visualized using Cytoscape version 3.8.2(29).

**Module-module interaction inference**

To examine possible module-module interactions across brain regions, Pearson correlations between module MEs were calculated among modules from all five brain regions in males and females, respectively. Module-module correlations with a p-value < 0.05 and |r| > 0.5 were used for visualization and the following analysis. Cytoscape 3.8.2 and sankeyNetwork function from the networkD3 R package were used for visualization.

**Associating WGCNA modules with human GWAS of cannabis use disorder**

To test whether THC-correlated or interconnected WGCNA modules were enriched in genes implicated in human GWAS of cannabis use disorder (CUD), we used the Mergeomics R package that integrates gene sets with GWAS summary statistics through functional genomics(26,30). GWAS summary statistics of a GWAS meta-analysis of CUD by Johnson et al.(31) from <https://www.med.unc.edu/pgc/download-results/> was used. SNPs were adjusted for linkage disequilibrium with a cutoff of 0.5 to remove redundant SNPs. GWAS SNPs were mapped to genes using the following mapping strategies: (1) genes within 20kb distance of SNPs and (2) genes under the regulation of tissue-specific expression and splicing quantitative trait loci (eQTLs/sQTLs) from GTEx v8 for brain regions matching or semi-matching those investigated for THC response. These include the amygdala, nucleus accumbens, frontal cortex, caudate and putamen (representing striatum), and substantia nigra (physically close to VTA). GWAS SNPs, their mapped genes, and the corresponding GWAS p values were then intersected with coexpression modules for each brain region using the Marker Set Enrichment Analysis (MSEA) function in Mergeomics to assess whether coexpression modules, compared to random gene sets, are enriched for genes with low p-value SNPs. Modules with an FDR < 0.05 are considered significantly enriched for GWAS-related genes.

Key driver genes from highlighted GWAS-associated modules were identified using wKDA and then visualized using Cytoscape 3.8.2 as described above.
