## Supplementary material for "Chronic adolescent exposure to cannabis in mice leads to sex-biased changes in gene expression networks across brain regions": Table S7

| **Key drivers of CUD & THC-associated modules** | | | | | |
| --- | --- | --- | --- | --- | --- |
|  | **DMS** | **Amy** | **NAc** | **PFC** | **VTA** |
| **Female** | GABRB2, PSD3, MEF2C, ATP2B1, NR2C2, ATF2, ADCY1, NFASC, TTYH2, SUN2, ERBB3, FA2H, NECTIN1 | FA2H, MAG, MAL, MOG, KLK6, GJB1, TMEM125, TF, CDC42EP1, OPALIN, RAB3A, LDOC1, ARHGDIG, TMEM130, C1QTNF4, RNF187, SYNGR3, BSCL2, SYP, ATP6V0D1 | **HAPLN4**, **KCNC1**, **ELAVL2**, SGSM1, ATP8A2, CLSTN3, **ZCCHC12**, RBFOX2, ANK1, SPTBN4 | JPH3, KCNH3, NEURL1, AGAP2, JPH4 | DDC, TH, SLC6A3, RET, SLC18A2, CHRNA6, KLHL1, FOXA2, FOXA1, SLC10A4, ZNF638, MATR3, RANBP6, PAIP2, ITGB8, SDCBP, PNPLA8, SEC23A, KPNA3 |
| **Male** | SYNGR1, WSCD2, FASN | ATP6V0A1, UBE2O, MADD, NISCH, DNM1, SMARCA4, CHD5, CELSR2, SPTBN2, MAPK8IP3 | RAB3A, **KCNC1**, **ELAVL2**, RAB3C, SNAP91, PNMAL2, **HAPLN4**, DNM1, SYT1, **ZCCHC12**, PJA2, WDR47, SRPK2, ITFG1, NCOA4, NAPB, NAPG, SUCLA2, YWHAH, CIT, PTPRS, CELSR3, PRRC2B, MECP2, CREBBP, MTUS2, BICDL1, PLAGL2, HCFC1, MLC1, RNF187 | SRPK2, G3BP2, FAM102B, RTN4, GLS, SEC23A, FAM49A, ITFG1, KIF3A, SH3BGRL | DNM1, APBA1, RBFOX2, SPTBN2, PI4KA, WNK2, CHRNB2, ARHGEF9, ATP6V0A1, ADD2 |
